# Genetic ablation of the mitochondrial ribosome in *Plasmodium falciparum* sensitizes the human malaria parasite to antimalarial drugs targeting mitochondrial functions

**DOI:** 10.1101/2020.01.14.906198

**Authors:** Liqin Ling, Maruthi Mulaka, Justin Munro, Swati Dass, Michael W. Mather, Michael K. Riscoe, Manuel Llinás, Jing Zhou, Hangjun Ke

## Abstract

The mitochondrion of malaria parasites contains clinically validated drug targets. Within *Plasmodium spp*., the mitochondrial DNA (mtDNA) is only 6 kb long, being the smallest mitochondrial genome among all eukaryotes. The mtDNA encodes only three proteins of the mitochondrial electron transport chain and ∼ 27 small, fragmented rRNA genes in length of 22-195 nucleotides. The rRNA fragments are thought to form a mitochondrial ribosome (mitoribosome), together with ribosomal proteins imported from the cytosol. The mitoribosome of *Plasmodium falciparum* has been shown to be essential for maintenance of the mitochondrial membrane potential and parasite viability. However, the role of mitoribosomes in sustaining the metabolic status of the parasite mitochondrion remains unknown. Here, among the 14 annotated mitoribosomal proteins of the small subunit of *P. falciparum*, we verified the localization and tested the essentiality of three candidates (PfmtRPS12, PfmtRPS17, PfmtRPS18), employing a CRISPR/Cas9 mediated conditional knockdown tool. Using immuno-electron microscopy, we provided evidence that the mitoribosome is closely associated with the mitochondrial inner membrane in the parasite. Upon knockdown of the mitoribosome, the parasites became hypersensitive to inhibitors targeting the *bc*_*1*_ complex, dihydroorotate dehydrogenase and *F*_*1*_*F*_*o*_ ATP synthase complex. Furthermore, knockdown of the mitoribosome blocked the pyrimidine biosynthesis pathway and reduced the pool of pyrimidine nucleotides. Together, our data suggest that disruption of the *P. falciparum* mitoribosome compromises the metabolic capability of the mitochondrion, rendering the parasite hypersensitive to a panel of inhibitors targeting mitochondrial functions.

## INTRODUCTION

Despite substantial success in reducing malaria morbidity and mortality over the last 15 years, malaria control has now encountered a serious bottleneck (1-3). Since 2015, statistics of global malaria illness (over 200 million cases per year) and death (∼ half a million per year) have remained almost static (1-3). In fact, malaria incidence had an increase of ∼ 5 million cases in 2016 alone (2), indicating a risk of resurgence of this devastating disease. To better combat malaria and potentially reach the goal of eradication, a deep understanding of the malaria parasite, a eukaryotic protozoan in the genus of *Plasmodium*, is urgently needed. Among the five *Plasmodium* species that infect humans, *Plasmodium falciparum* is responsible for more than 80% of the clinical cases and 90% of deaths. *P. falciparum* contains three distinct genomes, resident in the nucleus, the apicoplast (a relict non-photosynthetic plastid), and the mitochondrion, respectively (4). The spatial distribution of genetic material in the three subcellular compartments necessitates three distinct translation systems to translate local mRNAs to proteins. While the cytoplasmic ribosome resembles an 80S eukaryotic ribosome with some parasite-specific features (5,6), ribosomes in the organelles are of bacterial origin (7,8), which hold promise as novel antimalarial drug targets. The apicoplast ribosome has been long recognized as the target of several antibiotics (*e*.*g*., erythromycin and clindamycin) that exhibit a delayed death effect (9-11). However, the mitochondrial ribosome (mitoribosome) of malaria parasites has remained enigmatic.

In the mitochondria of eukaryotic cells, mitoribosomes translate genes encoded on the mitochondrial DNA (mtDNA), the majority of which are protein subunits of the mitochondrial electron transport chain (mtETC). Human mtDNA is a circular molecule of 16.6 kb, encoding 13 proteins, 22 tRNA genes, and 2 mitochondrial ribosomal RNA (rRNA) genes (12S and 16S) (12). However, in malaria parasites and other members of the phylum Apicomplexa, reductive evolution has likely driven the size of mtDNA to a minimum (13). Being an extreme case, *e*.*g*., the linear 6 kb mtDNA of malaria parasites is the smallest organellar genome among the entire Eukarya (14-16). It is highly compact, encoding only three proteins (cytochrome *b*, cytochrome *c* oxidase subunits I and III (*cox1, cox3*)), fragmented rRNA genes, but no tRNAs or other open reading frames (17). The three proteins comprise the highly hydrophobic cores of two proton pumps of the *Plasmodium* mtETC, Complex III (*bc*_*1*_ complex) and Complex IV (cytochrome *c* oxidase), with the estimated number of transmembrane helices present in the *cyt b, cox1*, and *cox3* subunits being 9, 13, and 7, respectively (TMHMM Server, v.2.0). The importance of mitochondrial protein translation in malaria parasites is underscored by the fact that the *bc*_*1*_ complex is a major antimalarial drug target. The clinical drug atovaquone, a hydroxynaphthoquinone, targets the Qo site of *bc*_*1*_ complex, substantially collapses the mitochondrial membrane potential (ΔΨm) and halts parasite growth (18-20). A long list of structurally distinct *bc*_*1*_ inhibitors are currently in the drug development pipeline (21), including the preclinical candidate, ELQ-300 (inhibitor of Qi site). ELQ-300 is highly potent and selective, long lasting, and effective against multiple lifecycle stages (22-24). Despite the significance of the mtETC, how the critical subunits are translated in the parasite mitochondrion is still poorly understood.

Arguably, the *Plasmodium* mitoribosome represents an unusually atypical ribosome due to its RNA composition consisting of many small and fragmented pieces (17). The 6 kb mtDNA encodes 27 rRNA “minigenes” annotated to be integrated into the small subunit (SSU, 12 rRNAs) and the large subunit (LSU, 15 rRNAs) of the mitoribosome (17). These rRNA fragments are sized between 22-195 nucleotides. Importantly, the short fragmented rRNA genes can be mapped to the core conserved structures of bacterial SSU and LSU rRNAs (16S and 23S) (17), suggesting their potential roles in protein synthesis. Fragmented mitoribosomal rRNAs have been observed in other free living single-celled organisms such as the ciliate *Tetrahymena thermophila* (6 fragmented rRNAs) (25) or the green alga *Chlamydomonas reinhardtii* (12 fragmented rRNAs) (26). However, the degree of rRNA fragmentation observed in the mitoribosome of malaria parasites greatly surpasses these examples. Recent ground-breaking advancements in cryo-EM technology have resolved the structures of mitoribosomes from yeast (27,28), mammals (29-34) and *Trypanosoma brucei* (a protist) (35). These studies have greatly increased our understanding of mitoribosomes and interactions of rRNA and ribosomal proteins (rProteins). However, all of these organisms contain long and continuous rRNAs in the mitoribosome, including *T. brucei*, which has relatively small rRNA genes (9S/620 nucleotides, 12S/1176 nucleotides) (35).

While knowledge of mitoribosomes composed of numerous fragmented rRNAs has remained limited, we recently verified the essentiality of the *P. falciparum* mitoribosome for the first time (36), using a CRISPR/Cas9-mediated conditional knockdown technique, the TetR-DOZI-aptamer system (37). We showed that knockdown of the conserved mitoribosomal LSU protein, PfmtRPL13 (PF3D7_0214200), caused a loss of ΔΨm, followed by parasite death (36). The essential nature of the mitoribosome in *Toxoplasma gondii*, another apicomplexan parasite, has also been reported (38). However, there is no literature on any of 14 annotated SSU proteins in *Plasmodium* (www.PlasmoDB.org). Here, we report data on three annotated SSU proteins, PfmtRPS12 (PF3D7_0412100), PfmtRPS17 (PF3D7_1365100) and PfmtRPS18 (PF3D7_1211500) in *P. falciparum*. In particular, genetic knockdown of PfmtRPS12 caused the parasites to become hypersensitive to a panel of antimalarial drugs that target mitochondrial functions.

## RESULTS

### A predicted map of the *Plasmodium* mitoribosome SSU

In *Plasmodium spp*., there are a total of 41 annotated mitoribosomal proteins (8), most of which are universally conserved proteins present in all ribosomes (www.PlasmoDB.org). The SSU of the *Plasmodium* mitoribosome is composed of 14 annotated proteins, at least 12 fragmented rRNAs and likely a number of additional ribosomal proteins that are not identifiable via currently available bioinformatics approaches. Based on the assembly map of the well-studied bacterial 30S ribosome (39) and the 6 kb sequence of *Plasmodium* mtDNA (17), we predicted relative positions of 12 mitoribosomal rRNA fragments in 5’ to 3’ directions according to their homology to different domains of the 16S rRNA molecule (Figure 1). The likely interactions of rRNA fragments and 14 annotated mitoribosomal proteins are also illustrated (Figure 1). Bacterial SSU proteins are classified into three categories according to the assembly hierarchy: primary (1°), secondary (2°) and tertiary (3°) (39). As shown in Figure 1, *Plasmodium* contains at least 3 out of 6 primary SSU proteins. Primary SSU proteins directly bind rRNA in the bacterial ribosome (39). *Plasmodium* also contains at least 4 out of 6 secondary SSU proteins and 4 out of 8 tertiary SSU proteins. In addition, 3 extra annotated SSU proteins in *Plasmodium* (S22, S29, and S35) have no bacterial homologs; hence, the positions of these three were not predicted. Recent work on *Toxoplasma gondii*, another apicomplexan parasite, showed that TgmS35 (TGME49_203620) is localized in the parasite mitochondrion and is essential for parasite growth (38). From the list of 14 *Plasmodium* SSU proteins, in this study, we chose to work with three putative proteins: primary (PfmtRPS17, PF3D7_1365100), secondary (PfmtRPS18, PF3D7_1211500) and tertiary (PfmtRPS12, PF3D7_0412100).

**Figure 1.**
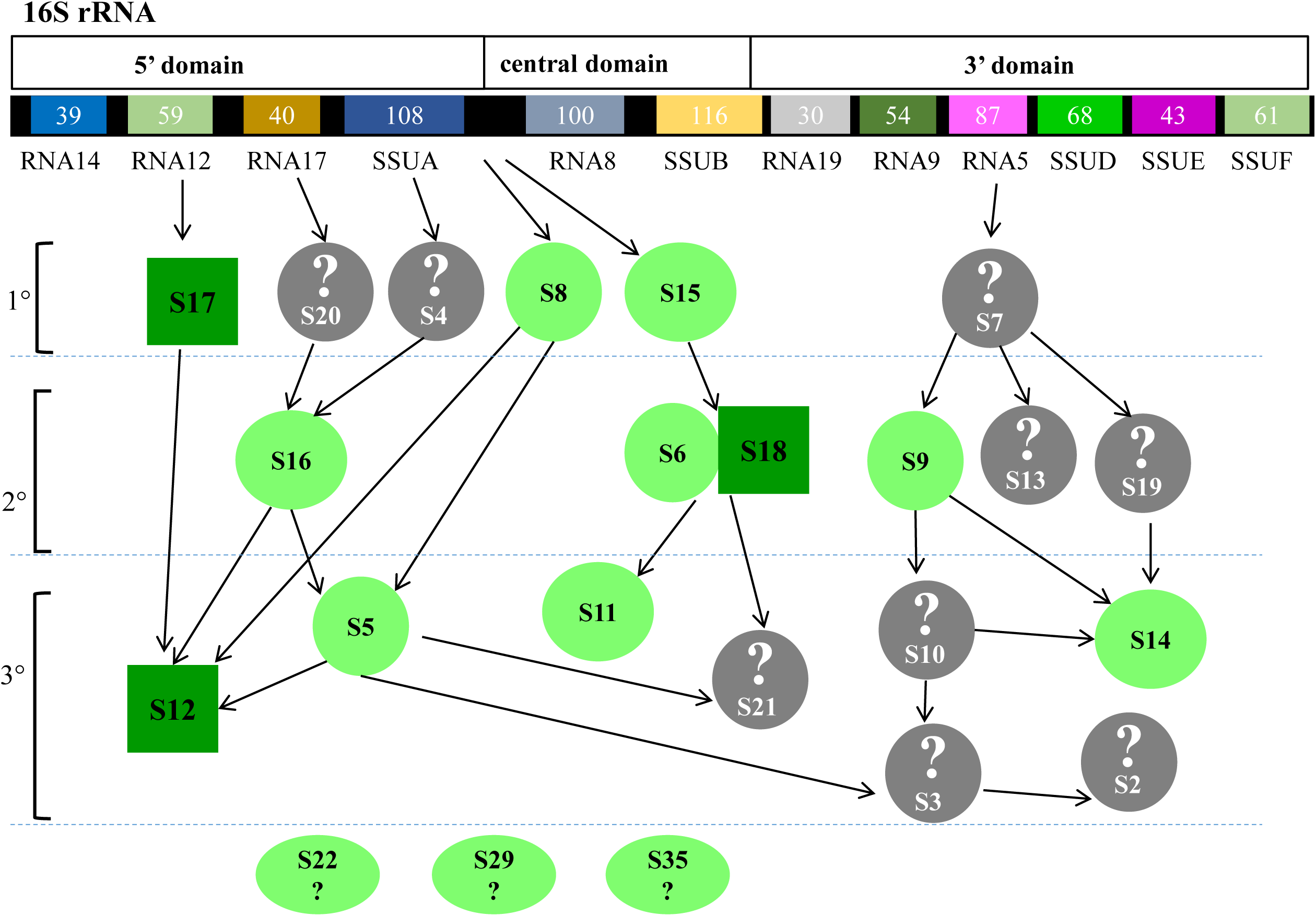
A map of the *Plasmodium* mitoribosome SSU. Based on the Nomura assembly map of the 30S ribosome, we highlight the estimated positions of the annotated rRNA fragments and proteins in the SSU of the *Plasmodium* mitoribosome. For each annotated rRNA fragment, its name and size is indicated. The rRNA fragments are mapped to 16S rRNA according to sequence homology. Fourteen annotated *Plasmodium* mitoribosomal proteins are highlighted in either dark and light green, which are classified into primary, secondary and tertiary proteins according to the Nomura assembly map. Interactions of rRNA and proteins observed in the 30S ribosome are shown in arrows. Three SSU proteins of the *Plasmodium* mitoribosome are chosen in this study (dark green squares). The bacterial SSU proteins that lack homologs in *Plasmodium* are highlighted in grey with a question mark. Three *Plasmodium* SSU proteins that lack homologous bacterial counterparts are highlighted in light green with a question mark. The positions of these three proteins (S22, S29, S35) are not predicted.

### PfmtRPS12, PfmtRPS17 and PfmtRPS18 are mitochondrial proteins

To determine if PfmtRPS12, PfmtRPS17 and PfmtRPS18 actually localize to the mitochondrion in *P. falciparum*, and if they are essential for parasite growth in asexual blood stages, we used a recently developed translational knockdown approach, the TetR-DOZI-aptamer system (37). Three transgenic parasite lines, D10-PfmtRPS12-3HA, D10-PfmtRPS17-3HA and D10-PfmtRPS18-3HA were generated from the D10 wildtype to individually tag and control expression of the three subunits (Experimental Procedures). As shown in Supplementary Figure 1A, the SSU genes were individually tagged with a triple HA tag. Meanwhile, the elements of the TetR-DOZI-aptamer system were also integrated at the genomic loci via the double crossover recombination approach facilitated by CRISPR/Cas9. As depicted in Supplementary Figure 1B, translation of the tagged mRNA was conditionally regulated by addition (ON) or removal (OFF) of the small molecule, anhydrotetracycline (aTc) (37). The genotype of each SSU transgenic line was confirmed by diagnostic PCR analysis (Figure 1C). The expression of the tagged SSU proteins was confirmed by Western blot, as shown in Figure 2A. The localization of each SSU protein was then examined by immunofluorescence assay (IFA); all three localized to the mitochondrion (Figure 2B).

**Figure 2.**
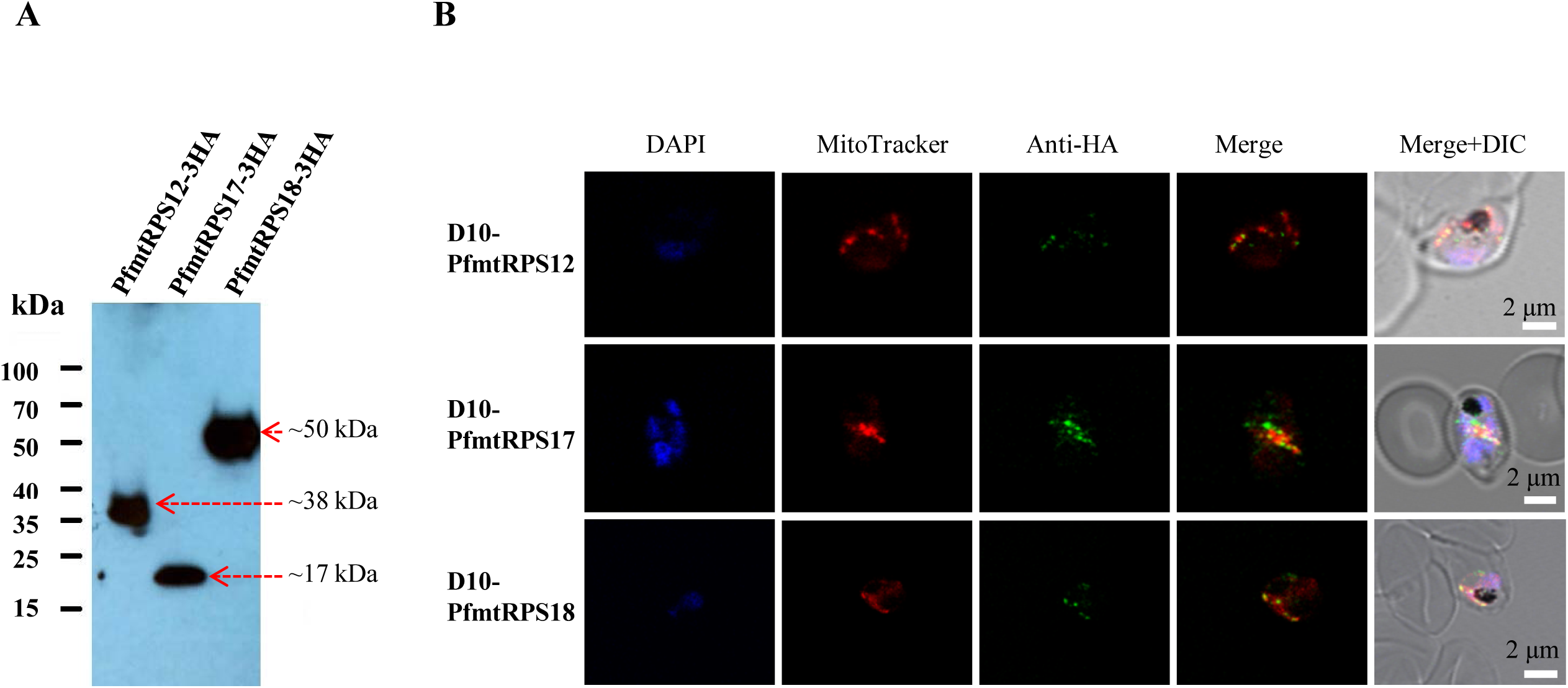
Expression and localization of PfmtRPS12, PfmtRPS17 and PfmtRPS18. (A) Detection of PfmtRPS12-3HA (∼ 38 kDa), PfmtRPS17-3HA (∼ 17 kDa), and PfmtRPS18-3HA (∼ 50 kDa) in transgenic parasites by Western blot. (B) PfmtRPS12-3HA, PfmtRPS17-3HA, or PfmtRPS18-3HA was localized to the parasite mitochondrion. A co-staining of anti-HA antibody and MitoTracker is shown. Note that the tubular mitochondrion in trophozoites often becomes artificially “fragmented” after fixation with paraformaldehyde.

### PfmtRPS18 appears to be located on the mitochondrial inner membrane

Our IFA data suggest that these three proteins are likely mitoribosomal proteins in the *Plasmodium* mitochondrion (Figure 2). Since mitoribosomes translate highly hydrophobic proteins of the mtETC, the exit tunnel is generally positioned toward the mitochondrial inner membrane (MIM) and directly contacts the MIM (40). This close interaction of mitoribosomes and MIM facilitates co-translational insertion of the multi-transmembrane protein products into the mtETC, which has been observed in mitochondria of several organisms (40). We used immuno-electron microscopy (immuno-EM) to verify if our tagged SSU proteins had a close proximity to the MIM. Since immuno-EM is technically more challenging than IFA, we chose to proceed with one SSU protein that has a relatively high expression level. A previous proteomic study that quantified expression profiles of over 2,000 proteins in the asexual blood stage of *P. falciparum* detected the expression level of PfmtRPS18 at the 75^th^ percentile relative to the levels found among the pool that contained trophozoite and schizont stage samples (41). The expression of PfmtRPS12 or PfmtRPS17, on the other hand, did not appear to be abundant enough for proteomic studies (41). Hence, we enriched D10-PfmtRPS18-3HA parasites using a magnetic column and performed immuno-EM studies (Experimental Procedures). Although gold particles sometimes were present at other sites, most of them were found in the mitochondria at or very near the MIM (Figure 3A and 3B), indicating that the PfmtRPS18 was likely associated with the MIM. As a negative control, omission of the primary antibody had a clean background (Figure 3C). Together, these data suggest the *Plasmodium* mitoribosome is likely bound to the mitochondrial inner membrane, as in other organisms.

**Figure 3.**
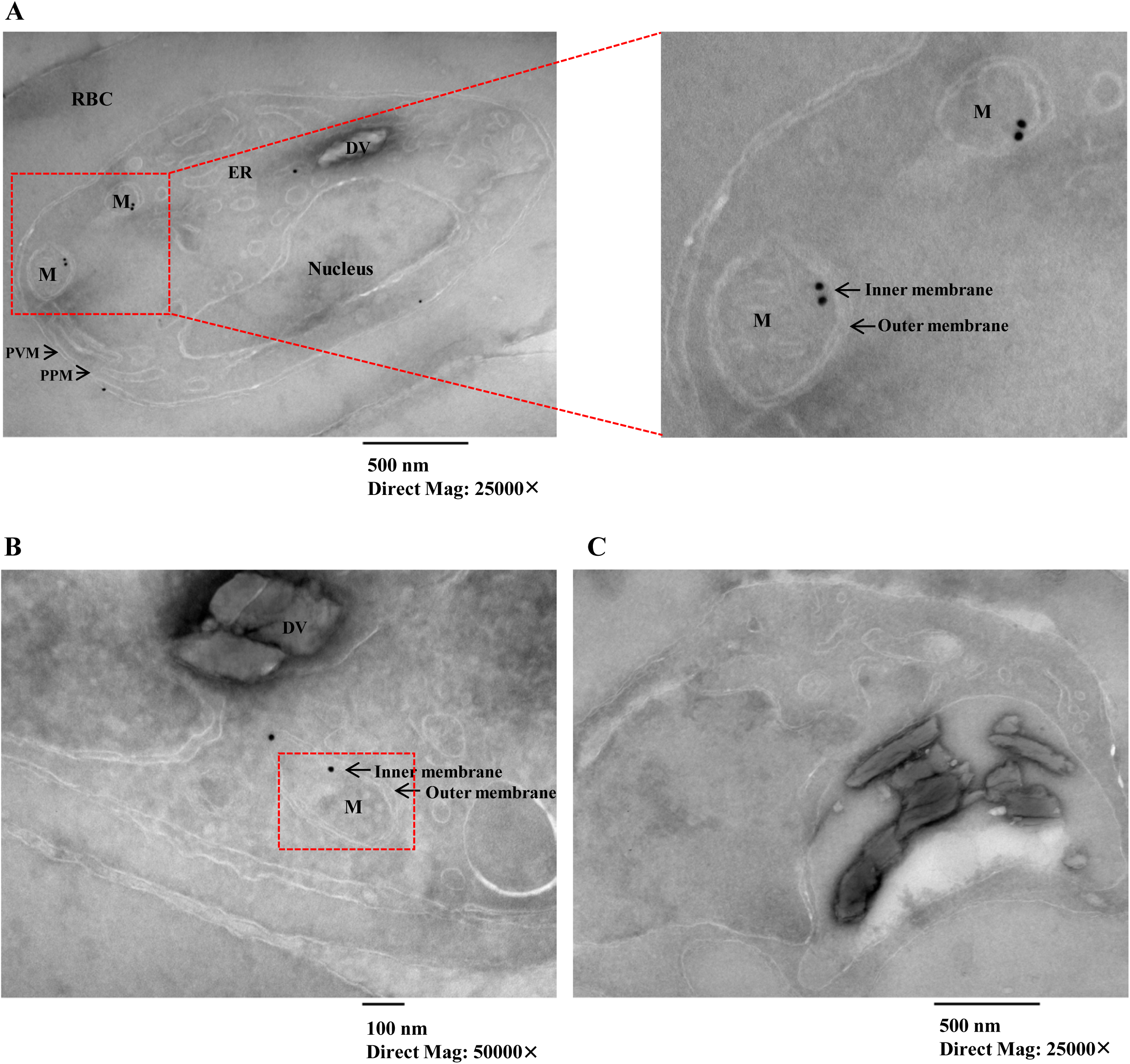
PfmtRPS18 is likely localized to the mitochondrial inner membrane. (A), (B) Images of D10-PfmtRPS18-3HA parasites. RBC, red blood cell. PVM, parasitophorous vacuole membrane. PPM, parasite plasma membrane. DV, digestive vacuole. ER, endoplasmic reticulum. M, mitochondrion. Black dots in red boxes indicate PfmtRPS18-3HA is localized to the mitochondrial inner membrane. (C) A control with omission of the primary antibody shows clear backgound. Mag, magnification.

### Loss of distinct SSU proteins leads to different growth phenotypes

Since these three SSU proteins likely occupy distinct sites in the mitoribosome (Figure 1), we wanted to know whether a genetic ablation of each individual protein would cause similar or different growth phenotypes. Using the TetR-DOZI-aptamer system (37), expression of three mitoribosomal proteins was regulated by aTc (Supplementary Figure 1B). Previously we found that 250 nM aTc itself did not interfere with growth of wildtype D10 parasites (36). The three SSU transgenic lines were maintained in aTc medium (250 nM) prior to knockdown assays. To individually assess the essentiality of three SSU subunits of the *P. falciparum* mitoribosome, we set up knockdown assays in the transgenic lines and monitored parasite growth over five intraerythrocytic cycles (IDCs) (Experimental Procedures). In particular, to minimize aTc carryover, the parasitized RBCs were isolated by a percoll gradient and exposed to medium with or without 250 nM aTc (Experimental Procedures). The knockdown efficiency was confirmed by Western blot, as shown in Supplementary Figure 2; expression of SSU proteins was substantially decreased in knockdown parasites after aTc removal.

We found that knockdown of different SSU proteins resulted in distinct growth defects. As shown in Figure 4B, knocking down PfmtRPS17 caused a severe growth arrest. Although the parasite appeared morphologically healthy under a light microscope after aTc had been removed for 1 IDC, detrimental changes were observed after 2 IDCs without aTc (Supplementary Figure 3B). After aTc removal for 3 IDCs or longer, D10-PfmtRPS17-3HA parasites were severely deteriorated. The essentiality of the PfmtRPS17 subunit is consistent with the fact that the bacterial S17 is a critical protein for assembly of the 30S ribosome. It directly binds to the 5’ end of 16S rRNA to stabilize the first step of SSU assembly (42). Knocking down PfmtRPS12, on the other hand, resulted in a mild growth arrest. As shown in Figure 4A and Supplementary Figure 3, in the first 2 IDCs without aTc, PfmtRPS12 parasites did not exhibit a loss in parasitemia or any detectable abnormality in morphology; however, the parasites started to show a decrease in growth rate in the 3^rd^ IDC (∼ 20% loss) and more noticeably in the 4^th^ IDC 4 (∼ 50% loss). After withdrawal of aTc for 4 or 5 IDCs, some PfmtRPS12 parasites displayed noticeable abnormalities in morphology (Supplementary Figure 3A). Despite these defects in growth rate and parasite morphology, PfmtRPS12 parasites were able to continuously grow after aTc removal for longer than 5 IDCs (data not shown). Previous studies in the 30S ribosome have shown that the bacterial S12 protein does not directly bind to 16S rRNA, but rather is involved in tRNA selection in the decoding center of the SSU (43). The mild phenotype seen upon knockdown of PfmtRPS12 suggests that this *Plasmodium* homolog has a significant but perhaps not critical function in the mitoribosomal complex. In D10-PfmtRPS18-3HA parasites, removal of aTc had negligible effect on both parasite growth and morphology (Figure 4C and Supplementary Figure 3C), suggesting that PfmtRPS18 is not essential for asexual growth *in vitro*, consistent with the results from the recent large-scale piggyBac transposon mutagenesis survey, in which PfmtRPS18 was found to be mutable (44). In all, our data show that knocking down different SSU proteins affects the parasites, and presumably the mitoribosome, to varying degrees.

**Figure 4.**
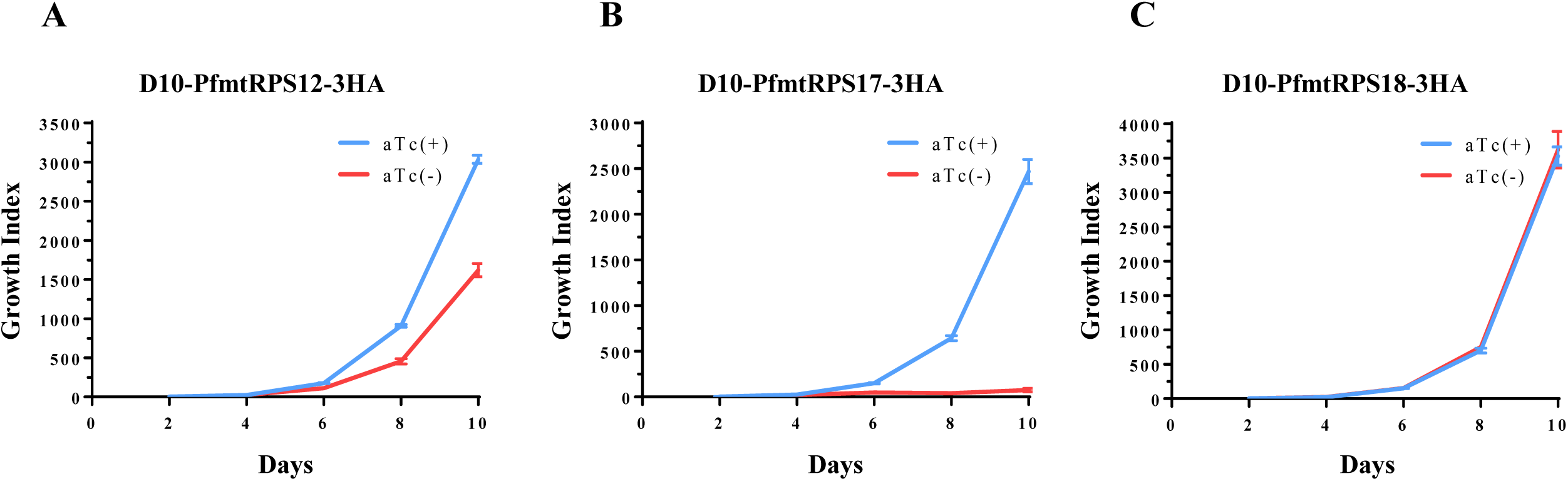
Effect on parasite survival upon individual knockdown of PfmtRPS12, PfmtRPS17 and PfmtRPS18. Representative growth curves of D10-PfmtRPS12-3HA (A), D10-PfmtRPS17-3HA (B) or D10-PfmtRPS18-3HA (C) grown in aTc^+^ (blue) and aTc^-^ (red) conditions. Growth index is the cumulative parasitemia which is the multiplication of parasitemia and split factors (1:5 split every IDC) over the time course. Data shown are the mean ± s.d. of three biological replicates (n = 3). aTc, anhydrotetracycline.

### Loss of distinct SSU proteins results in a decrease in *bc*_*1*_ enzymatic activity

In *Plasmodium*, the mitoribosome solely translates three proteins that are incorporated into the *bc*_*1*_ complex (*cyt b*) and Complex IV (*cox1* and *cox3*). Due to lack of antibodies against these proteins, a direct measurement of these protein products in our knockdown parasites is not feasible at present. We have previously established a protocol to measure the *bc*_*1*_ enzymatic activity in extracts of parasites grown *in* vitro (36), which in turn reflects the efficiency of mitochondrial protein translation. Hence, we isolated mitochondria from our SSU transgenic parasites grown in large culture volumes with and without aTc and assayed the *bc*_*1*_ activity in each mitochondrial preparation (Experimental Procedures). Since knockdown of individual SSU proteins led to distinct growth defects (Figure 4), mitochondria were harvested after a different number of IDCs of growth following aTc removal. For example, mitochondria from D10-PfmtRPS17-3HA parasites were isolated after aTc removal for 2 IDCs, at which point the parasites had started to show defects but the parasitemia did not declined too dramatically. Likewise, mitochondria from D10-PfmtRPS12-3HA parasites were harvested at the 4th IDC after withdrawal of aTc when they showed moderate growth defects. Since PfmtRPS18 knockdown produced no detectable growth defects, mitochondria from D10-PfmtRPS18-3HA parasites were isolated in cultures maintained without aTc for 5 IDCs. As controls, mitochondria from D10-PfmtRPS12-3HA, D10-PfmtRPS17-3HA, and D10-PfmtRPS18-3HA parasites were isolated in cultures grown continuously in the presence of aTc.

As shown in Figure 5, in all three SSU knockdown parasite lines, the enzymatic activity of *bc*_*1*_ was significantly compromised. In comparison to that of the controls, the *bc*_*1*_ enzymatic ability was reduced by 52% in D10-PfmtRPS12-3HA knockdown parasites, 53% in D10-PfmtRPS17-3HA knockdown parasites and 30% in D10-PfmtRPS18-3HA knockdown parasites. As a control, 1 µM atovaquone completely diminished the *bc*_*1*_ enzymatic activity, validating the assay. While all three knockdown lines lost significant *bc*_*1*_ enzymatic activity, knockdown of PfmtRPS18 had a milder effect on *bc*_*1*_ than that of PfmtRPS12 or PfmtRPS17. This is consistent with the non-essential role of PfmtRPS18 in parasite survival as shown in Figure 4C. However, it’s interesting to note that PfmtRPS18 parasites can tolerate a 30% loss in *bc*_*1*_ activity without suffering any noticeable morphological changes or growth defects. In both PfmtRPS12 and PfmtRPS17 lines, the effect of knockdown on the *bc*_*1*_ enzymatic activity was similar; however, a longer period of aTc removal was needed in PfmtRPS12 (4 IDCs) as compared to PfmtRPS17 (2 IDCs) to reduce *bc*_*1*_ activity by > 50%. Overall, as expected, these data clearly show knockdown of SSU proteins affects the mitochondrial protein translation and reduces the *bc*_*1*_ activity.

**Figure 5.**
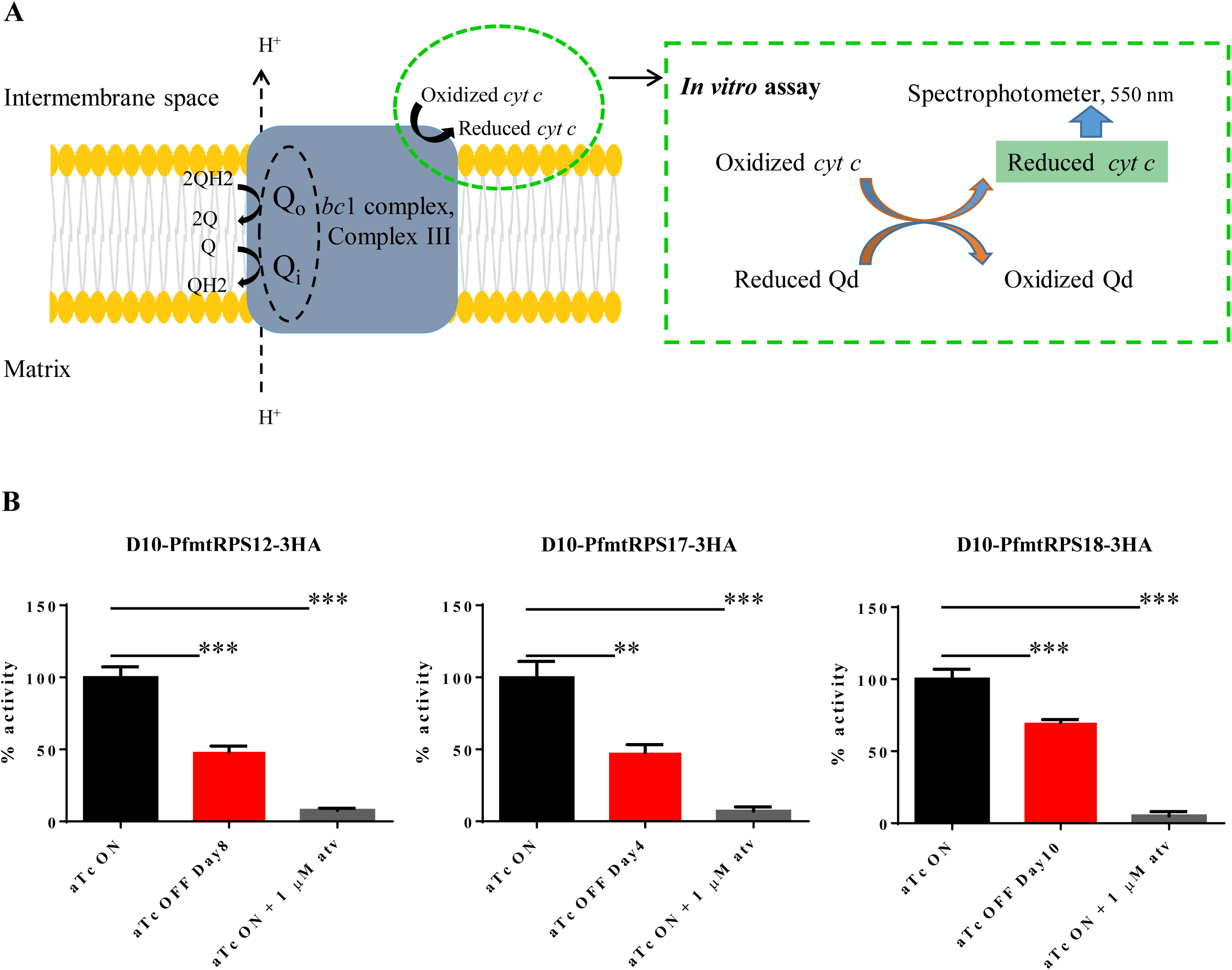
Knockdown of PfmtRPS12, PfmtRPS17 and PfmtRPS18 leads to a reduced *bc*_1_ activity. (A) A schematic representation of the Q cycle in the *bc*_1_ complex. *In vitro*, the rate of reduction of oxidized *cyt c* by reduced Qd catalyzed by the *bc*_1_ complex was measured by a light-scatter rejecting UV/VIS spectrometer under 550 nm. B, The *cyt c* reduction activity of the *bc*_1_ complex is reduced in all three SSU knockdown lines, (A) D10-PfmtRPS12-3HA (B) D10-PfmtRPS17-3HA and (C) D10-PfmtRPS18-3HA. Atv (atovaquone) was used at 1 µM to completely abolish the *bc*_1_ activity. Error bars (s.e, standard error) and statistical analysis (Student’s t test) are derived from three biological replicates (n = 3). **, *p* < 0.01; ***, *p* < 0.001.

### Loss of PfmtRPS12 causes a hypersensitivity to mtETC-targeting drugs

The mitochondrion of *Plasmodium* has been successfully exploited as a major antimalarial drug target (21). The *bc*_*1*_ complex is the target of the clinical drug atovaquone, the preclinical candidate ELQ-300 and other inhibitors in the development pipeline (21). The *de novo* pyrimidine biosynthetic enzyme, dihydroorotate dehydrogenase (DHOD), is the target of DSM-265, a candidate currently undergoing a Phase IIa clinical trial (45). Complex V (*F*_*1*_*F*_*o*_ ATP synthase) is believed to be the target of proguanil (46), the second compound in the registered drug Malarone (a combination of atovaquone and proguanil). All of these drugs target mtETC-related proteins (Figure 6A). We investigated whether disruption of the *Plasmodium* mitoribosome would have a synergistic effect on the action of antimalarial drugs that inhibit enzymes of the mtETC. In our previous report (36), the PfmtRPL13 knockdown line was generated using the pre-gRNA construct carrying Cas9 and the yeast DHOD (yDHOD). Previous studies revealed that heterologous expression of yDHOD in *P. falciparum* rendered the parasite insensitive to atovaquone and other mtETC drugs (46); thus, the PfmtRPL13 line was not suitable for testing drugs related to mitochondrial functions. Here, we generated the three SSU knockdown lines using a pre-gRNA construct lack of yDHOD (Experimental Procedures), which are suitable for drug testing.

**Figure 6.**
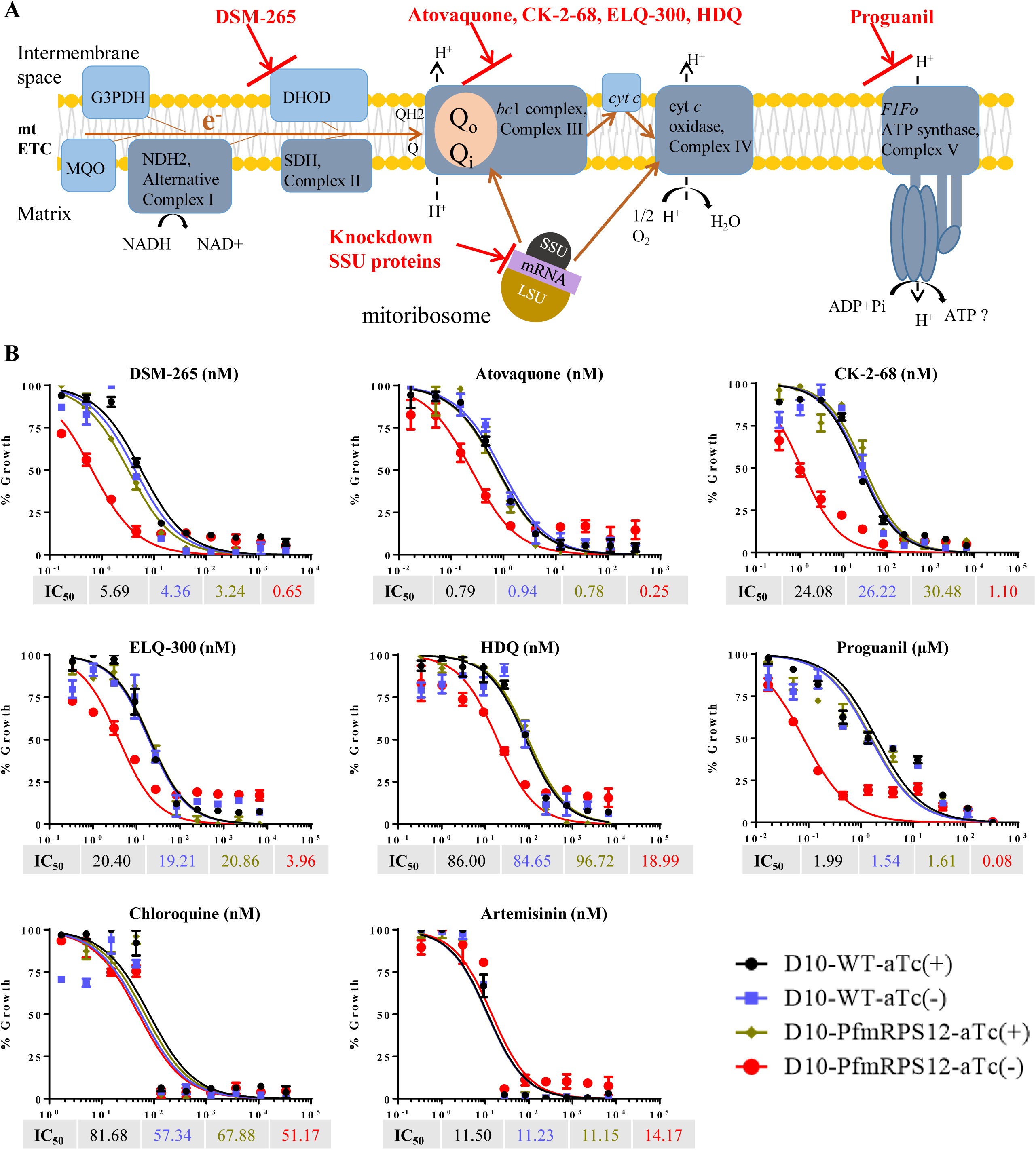
Knockdown PfmtRPS12 renders the parasite become hypersensitive to inhibitors targeting mitochondrial functions. (A) The *Plasmodium* mtETC and *F*_*1*_*F*_*o*_ ATP synthase complex. G3PDH, glycerol-3-phosphate dehydrogenase. MQO, malate-quinone oxidoreductase. NDH2, ubiquinone oxidoreductase. DHOD, dihydroorotate dehydrogenase, the target of DSM-265. SDH, succinate dehydrogenase. Cyt c, soluble cytochrome *c*. Q, ubiquinone. QH2, ubiquinol. *bc*_1_ complex Q_o_ site, the target of atovaquone, CK-2-68. *bc*_1_ complex Q_i_ site, the target of ELQ-300, HDQ. Complex V, *F*_*1*_*F*_*o*_ ATP synthase, the target of proguanil. (B) Hypersensitivity to DSM-265, atovaquone, CK-2-68, ELQ-300, HDQ, and proguanil in PfmtRPS12 knockdown parasites. Data shown are the mean ± s.d. of duplicates and are representative of n = 3 independent experiments. Chloroquine and artemisinin served as controls.

We performed standard 3-day SYBR Green I drug inhibition assays beginning with ring stage parasites in the presence or absence of aTc. Since removal of aTc affected the SSU lines distinctly (Figure 4), we started drug assays after prior removal of aTc for 1.5 IDCs with the D10-PfmtRPS17-3HA line, 2.5 IDCs with the D10-PfmtRPS12-3HA line, and 4.5 IDCs with the D10-PfmtRPS18-3HA line. For instance, in the case of the PfmtRPS12 line, the drug assay covered days of 5-8 post aTc removal, which is when the parasites begin to exhibit a measurable growth defect. As shown in Figure 6B, upon removal of aTc, PfmtRPS12 parasites displayed hypersensitivity to DSM-265, atovaquone, CK-2-68 (a *bc*_*1*_ inhibitor) (47), ELQ-300, HDQ (a *bc*_*1*_ inhibitor) (48), and proguanil. As controls, these parasites did not exhibit hypersensitivity to drugs that do not target the mtETC or related mitochondrial processes, *e*.*g*., chloroquine and artemisinin. Hence, the hypersensitivity to DSM-265, proguanil and other mtETC-targeting drugs observed in D10-PfmtRPS12-3HA parasites was specific to aTc removal, not likely due to a secondary non-specific effect. In addition, D10 wildtype parasites were also tested with the same panel of drugs in the presence and absence of aTc, and they did not show any differences in drug sensitivity (Figure 6). Together, these results provide strong evidence that genetic ablation of the *Plasmodium* mitoribosome sensitizes the parasite to antimalarial drugs targeting the mtETC and Complex V.

Interestingly, however, this synthetic lethality only occurred in the PfmtRPS12 line, in which a genetic knockdown caused a significant fitness cost but did not severely affect parasite growth. In PfmtRPS18 parasites, they responded to all the tested drugs in a similar manner with or without aTc (Supplementary Figure 4). This was likely due to the fact that knockdown of PfmtRPS18 did not result in any defects in parasite growth (Figure 4 and Supplementary Figure 3C). On the other hand, knockdown of PfmtRPS17 had a dramatic effect by itself. When combined with antimalarial drugs, PfmtRPS17 knockdown parasites exhibited hypersensitivity to all the drugs tested in a non-specific manner, including DSM-265, atovaquone, CK-2-68, ELQ-300, HDQ, proguanil, chloroquine and artemisinin (Supplementary Figure 4) (see Discussion).

### Metabolic profiling upon knockdown of PfmtRPS17

In the data described above, we observed that genetic knockdown of SSU proteins in the *Plasmodium* mitoribosome resulted in parasite growth arrest (Figure 4, PfmtRPS12/ PfmtRPS17), reduction of *bc*_*1*_ activity (Figure 5, all lines), and synthetic lethality with antimalarials targeting the mtETC (Figure 6, PfmtRPS12). Since in asexual blood stages the parasite mitochondrion does not synthesize a significant amount of ATP via oxidative phosphorylation, but primarily acts as a metabolic hub, we hypothesized that knocking down the mitoribosome could perturb the metabolic stability of the parasite mitochondrion. To address this, we chose to work with the D10-PfmtRPS17-3HA line, because severe growth arrest of this line was triggered in a short period of time upon removal of aTc. As shown above (Figure 4 and Supplementary Figure 3), in the absence of aTc, although PfmtRPS17 parasites showed no noticeable growth arrest in the 1^st^ IDC (48 h), they displayed a reduced parasitemia and abnormal morphologies in the 2^nd^ IDC (96 h). Therefore, we set up the knockdown experiment and extracted metabolites for metabolomic analysis every 12 h between 24 and 96 h post aTc removal. Hence, samples at 7 timepoints were collected from both cultures grown with and without aTc (Experimental Procedures). This time course covered the time period before and after growth arrest of PfmtRPS17 parasites took place. However, it did not include samples of IDCs 3 and 4 when parasites were too severely ill and a metabolic perturbation could be a secondary effect of parasite death. As a positive control, PfmtRPS17 parasites grown in the presence of aTc were treated with 10 nM atovaquone for 4 h, and their metabolites were extracted and analyzed (Experimental Procedures).

As shown in Supplementary Figure 5, we detected a total of 104 metabolites in the time course samples, including metabolic intermediates of nucleotide metabolism, TCA cycle, glycolysis, amino acid metabolism, pyrimidine biosynthesis and others. In the first three timepoints (24, 36, 48 h), knocking down PfmtRPS17 led to fluctuations in the levels of many metabolites, but none decreased or increased in abundance by more than 2-fold in the absence of aTc. This data again indicates that PfmtRPS17 parasites were largely healthy upon removal of aTc for 1 IDC. However, beyond the timepoint of 60 h, the levels of the pyrimidine *de novo* synthesis pathway intermediates, N-carbamoyl-L-aspartate and dihydroorotate, increased significantly in the knockdown parasites (Figure 7). As a positive control, atovaquone treatment in PfmtRPS17 parasites grown under aTc also resulted in accumulations of these two intermediates. As shown previously (49), blockage of pyrimidine biosynthesis is a classical metabolic signature of antimalarial drugs targeting the parasite mtETC, because a direct chemical inhibition of mtETC subsequently blocks the activity of parasite DHOD (46). On the other hand, from the timepoints of 84 h and beyond, ten metabolites in the pyrimidine nucleotide pool and TCA cycle decreased in abundances by more than 2-fold in the absence of aTc (Metabolites 1-10, Supplementary Figure 5). In atovaquone treated samples, three of these metabolites (dTDP, CDP, citrate/isocitrate) also displayed > 2-fold decrease in their relative abundances. Overall, our data show knockdown of the *Plasmodium* mitoribosome largely recapitulates the effect of atovaquone on inhibiting pyrimidine biosynthesis.

**Figure 7.**
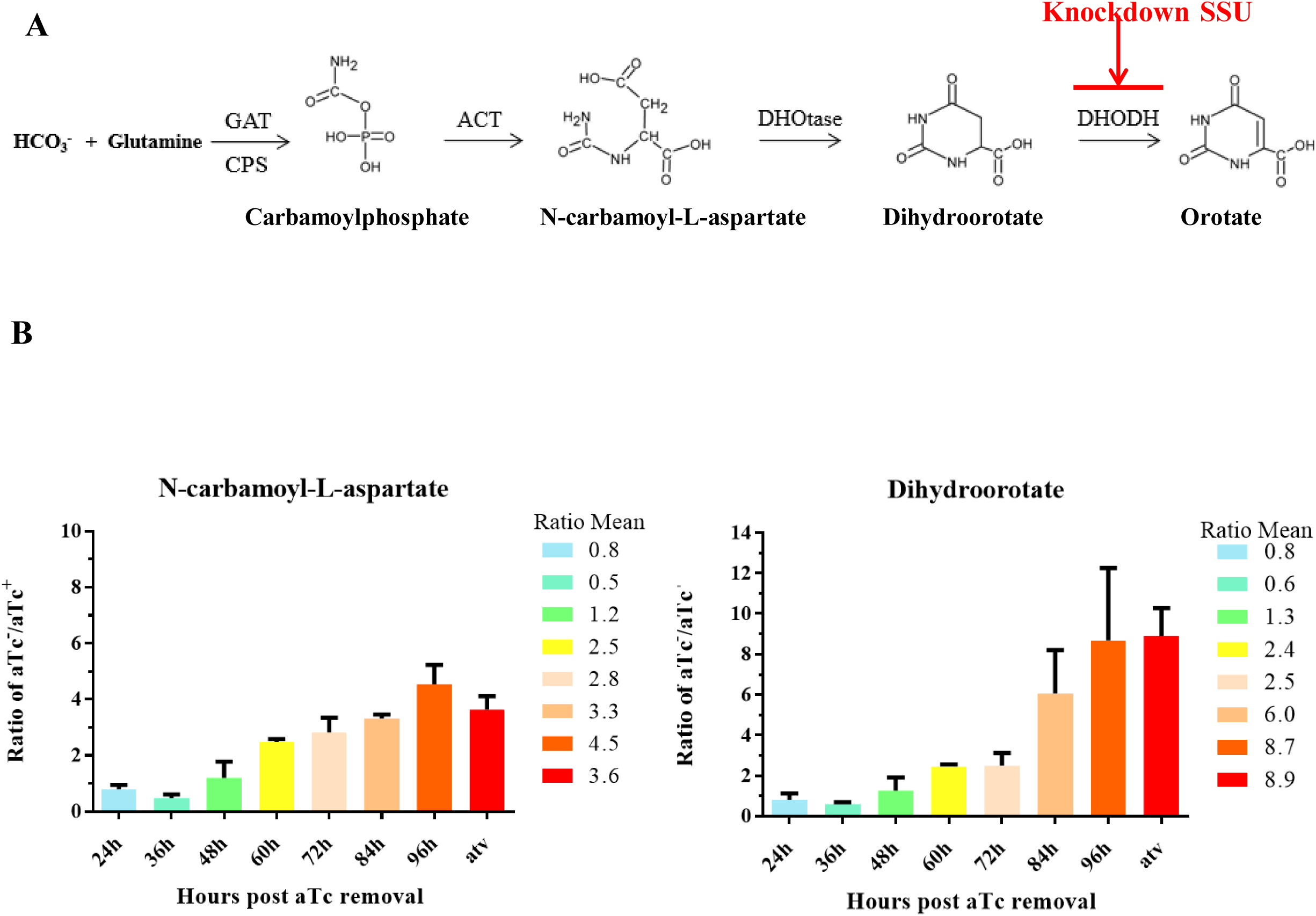
Loss of PfmtRPS17 results in accumulation of pyrimidine biosynthesis precursors. The *P. falciparum de novo* pyrimidine biosynthesis pathway. GAT, glutamine amidotransferase. CPS, carbamoyl phosphate synthase. ACT, aspartate carbamoyl transferase. DHOtase, dihydroorotase. DHODH, dihydroorotate dehydrogenase. Knockdown SSU denotes the blockage in conversion of dihydroorotate to orotate following disruption of the mitochondrial electron transport as shown in Figure 6. (B) Metabolic abundance of N-carbamoyl-L-aspartate and Dihydroorotate in aTc^-^ compared to aTc^+^ conditions. y axis, ratio of the metabolite in aTc^-^ compared to aTc^+^; x axis, timepoints post removal of aTc. The mean ratio of the metabolite at different timpoints was shown. Data shown in mean+s.e. were derived from three biological replicates (n = 3). Statistical analysis was done by one-way ANOVA test (N-carbamoyl-L-Aspartate, *p* < 0.01; Dihydroorotate, *p* < 0.05). Atv (atovaquone) was used at 10 nM in PfmtRPS17 parasites grown under aTc for 4 hrs.

## DISCUSSION

With the exception of one anaerobic protist that has lost its mitochondrion-related organelle (50), all eukaryotic organisms contain mitochondria or mitochondrially-derived organelles (*e*.*g*., mitosomes). In model systems like yeast and human cells, the mitochondrion is often considered the “power house” of the cell; however, ATP synthesis via oxidative phosphorylation is not always a major contribution of the mitochondrion in some deeply divergent parasitic protozoans (51). For example, in the blood stage infection of *Plasmodium spp*., the parasite mitochondrion mainly serves as a metabolic hub to provide essential metabolites, *e*.*g*., pyrimidine precursors, ubiquinone, iron-sulfur clusters and more. To fulfill these metabolic roles, a functional mtETC is needed to pump protons across the mitochondrial inner membrane (MIM) to build up a proton electrochemical gradient, with the mitochondrial membrane potential, ΔΨm, as its major component. The energy saved in ΔΨm is then utilized by the mitochondrion to import proteins and metabolic precursors from the cytosol. Although many mtETC subunit proteins are nuclearly encoded in apicomplexan parasites, as in other eukaryotes, three highly hydrophobic transmembrane proteins (*cyt b, cox1*, and *cox3*) have been evolutionarily retained on the mtDNA and are translated inside the mitochondrion. Despite the significance of mtETC in apicomplexan parasites, few studies have been done to understand mechanisms of mitochondrial protein translation. It was only recently demonstrated that the mitoribosome is absolutely essential in these parasites (36,38). In addition, we found that mitochondrial protein translation is not only critical for maintenance of mitochondrial functions, but is also important to keep the integrity of the growing MIM as the parasite matures (36). As shown previously, we found that the mitoribosome is still needed when a bypass of mtETC is provided by the expression of yDHOD (36). In this work, we performed genetic, biochemical and functional analyses to further study the mitoribosome in *Plasmodium falciparum*.

### What can we learn about the Plasmodium mitoribosome from genetic studies?

Similar to what we previously reported using an LSU tagged line (PfmtRPL13) (36), knockdown of SSU proteins in the *Plasmodium* mitoribosome resulted in a series of lethal events, including reduction of *bc*_*1*_ activity (Figure 5), collapse of parasite morphology (Supplementary Figure 3), and loss of parasitemia (Figure 4). However, our data also yield important insights into the distinct roles of SSU proteins in the *Plasmodium* mitoribosome. As shown in Figure 4, knockdown of each of the three SSU proteins affected the mitoribosome to varying degrees from negligible (PfmtRPS18) to mild (PfmtRPS12) and severe (PfmtRPS17). The non-essential nature of PfmtRPS18 was also reported in a recent comprehensive piggyBac-insertion mutagenesis screen (44), although the homolog of the rodent malaria parasite *P. berghei* appears to be essential (52). S12 is essential in both *P. berghei* (52) and *P. falciparum* (44), whereas S17 was found to be essential in *P. falciparum* (44) (not assessed in *P. berghei* (52)). We reason that the position of a particular SSU protein inside the mitoribosomal complex is critical for its function. Unfortunately, to date, no structure of a mitoribosome or ribosome comprised of many fragmented rRNAs is available. Hence, it is difficult to explain why not all SSU proteins of the *Plasmodium* mitoribosome are critical. As shown in Figure 1, we mapped the available rRNA fragments and 14 SSU proteins of the *Plasmodium* mitoribosome to the bacterial 30S ribosome. In the bacterial ribosome, S17 is one of the earliest proteins to bind the 5’ end of 16S rRNA and directly interacts with helices H7 and H11(42). Although PfmtRPS17 has relatively low sequence similarity with the bacterial homolog (27% identity), a predicted structure of PfmtRPS17 by I-Tasser (53) is superimposable on the crystal structure of bacterial S17 (data not shown). Moreover, RNA12 in the *Plasmodium* mitoribosome is homologous to the region of 16S rRNA where helices H7 and H11 are located (13,17). These considerations combined with our data suggest that the absolute necessity of PfmtRPS17 likely devolves from direct binding of the protein to rRNA (likely, RNA12). Whether PfmtRPS12 or PfmtRPS18 could directly bind to rRNA remains unknown (or unpredictable) at present.

### How is a mitoribosome attached to the mitochondrial inner membrane?

For the first time, we have demonstrated a close association of the mitoribosome to the MIM in an apicomplexan parasite (Figure 3). Compared to other ribosomes, a unique feature of the mitoribosome is that all or most of its protein translation products are highly hydrophobic mtETC subunits. Studies of mammalian mitoribosomes with cryo-electron tomography have clearly shown the physical attachment of the mitoribosome to the MIM (40). In yeast and mammals, an increasing list of factors has been found that participate in mediating the interaction of the mitoribosome and the MIM in a structure called the mitoribosomal interactome. For example, the mitoribosomal protein MrpL45 (40) (UniProtKB-Q9BRJ2) and the MIM receptors (54), Mba1 (YBR185C) and Mrx15 (YNR040W), are critical subunits of the mitoribosomal interactome. However, a homology search using sequences of these proteins in PlasmoDB failed to identify any homologous proteins in *Plasmodium*, suggesting that the molecular partners of the mitoribosomal interactome in malaria parasites are highly divergent, and, if identified, they may be exploited to develop selective antimalarial inhibitors.

### Is the Plasmodium mitoribosome a good antimalarial drug target?

Previous studies have validated two antimalarial drug targets, *bc*_*1*_ and DHOD, in the *Plasmodium* mitochondrion. Inhibition of *bc*_*1*_ blocks the recycling of ubiquinone and in turn shuts down ubiquinone-dependent enzymes in the mtETC, including DHOD, which is essential in the pyrimidine biosynthesis pathway (46). Disruption of the *Plasmodium* mitoribosome simultaneously diminishes core components of the mtETC and DHOD. As shown in Figure 7 and Supplementary Figure 5, knockdown of the mitoribosomal SSU (in D10-PfmtRPS17-3HA line) phenocopied the metabolic effect of atovaquone on the parasite. In particular, 60-96 h post aTc removal, we observed an accumulation of pyrimidine biosynthesis intermediates, N-carbamoyl-L-aspartate and dihydroorotate, and a concurrent decrease in pyrimidine nucleotides (thymine/cytosine/uracil mono-/di-phosphate). Importantly, these metabolic changes emerged at a point when the parasites started to display noticeable growth defects, suggesting that a metabolic blockage of the pyrimidine biosynthetic pathway was primarily responsible for parasite death. Although we did not test metabolic profiles in the parasites when aTc was removed for longer than 2 IDCs, we expect continued knockdown of PfmtRPS17 would further reduce the pyrimidine pool and cause other abnormalities leading to parasite death.

Besides DHOD and pyrimidine metabolism, ablation of mitochondrial protein translation would also abolish other essential metabolic pathways of the mitochondrion. One of those is iron-sulfur (Fe-S) cluster biosynthesis. Fe-S cluster containing proteins are widely distributed in the nucleus and cytosol and are involved in numerous biochemical pathways, such as electron transfer, catalysis, DNA repair, and ribosome biogenesis (55,56). As in other eukaryotes, the mitochondrial Fe-S cluster biogenesis pathway in *Plasmodium* provides these redox centers to proteins in extra-mitochondrial compartments, as well as the mitochondrion. Indeed, the most conserved function of mitochondria or mitochondrially derived organelles is synthesis of iron sulfur clusters. For instance *Cryptosporidium*, another apicomplexan parasite, maintains several mitosomes per cell to accommodate Fe-S biosynthesis even though this genus lacks mtDNA (57). Malaria parasites contain dozens of proteins needed to perform Fe-S cluster biogenesis in the mitochondrion (4). However, these proteins are generally very similar to the counterparts in yeast and humans; thus, the likelihood of selectively inhibiting the parasite’s mitochondrial Fe-S cluster biogenesis is potentially low. As shown previously, disruption of the *Plasmodium* mitoribosome collapses ΔΨm (36). Although not directly tested, we speculate that genetic ablation of the *Plasmodium* mitoribosome subsequently blocks the mitochondrion’s ability to synthesize Fe-S clusters, due to the reduced ΔΨm, which hinders import of the enzymes involved in Fe-S cluster biogenesis, and contributes to parasite demise.

Further, we found that genetic knockdown of the *Plasmodium* mitoribosome had a synergistic effect with antimalarial drugs targeting the mitochondrion in the D10-PfmtRPS12-3HA line, in which a genetic ablation caused a significant but not severe growth failure (Figure 6). As shown in Supplementary Figure 4, knockdown of PfmtRPS18 did not affect parasite fitness, so the parasites responded to drugs in a similar way regardless of the growth condition (with or without aTc). However, with the D10-PfmtRPS17-3HA line, knockdown itself was severely detrimental, and increasing concentrations of drug had a minor role. Thus, the parasite became hypersensitive to antimalarials targeting both mitochondrial and extra-mitochondrial pathways (Supplementary Figure 4). Together, our studies suggest that the *P. falciparum* mitoribosome is a good antimalarial drug target. We predict chemical disruption of the *Plasmodium* mitoribosome, if suitable inhibitor compounds were to be developed, would simultaneously impact multiple essential pathways in the parasite mitochondrion. Hence, future inhibitors targeting the *Plasmodium* mitoribosome should be good partner drugs in combination with antimalarials targeting either the mitochondrion or other cellular pathways.

## EXPERIMENTAL PROCEDURES

### 1. Plasmid construction

The pAll-In-One (pAIO) vector was kindly provided by Dr. Joshua Beck at Iowa State University (58). This plasmid bears Cas9 from *Streptococcus pyogenes* and a guide RNA (gRNA) expression cassette. The Cas9 CDS is flanked by an yDHOD (yeast dihydroorotate dehydrogenase) gene on the 5’ end and a Flag tag on the 3’ end. We previously removed the yDHOD gene from pAIO, yielding a pre-gRNA construct without yDHOD designated pAIO-Cas9-yDHOD(-) (48). In pAIO-Cas9-yDHOD(-), we performed a series of cloning steps to replace *BtgZ*I with *EcoR*I for cloning gRNAs and modified the trans-activating sequence of the gRNA according to published studies (59,60) to enhance its binding to Cas9 (details in the Supporting Information). These procedures yielded a new pre-gRNA construct named M-Cas9-yDHOD(-). In M-Cas9-yDHOD(-), we removed the Flag tag at the C-terminus of Cas9, yielding another construct named NF-Cas9-yDHOD(-). All gRNAs used in this study were selected based on analysis by the Eukaryotic Pathogen CRISPR guide RNA Design Tool (http://grna.ctegd.uga.edu/). gRNAs were individually cloned into NF-Cas9-yDHOD(-) using DNA assembly reactions (NEBuilder HiFi DNA Assembly Master Mix, New England Biolabs®, Inc.). The TetR-DOZI-aptamer conditional knockdown system was kindly provided by Dr. Jacquin Nile’s group at MIT (37). From the original pMG75-ATP4 vector bearing the TetR-DOZI and aptamer elements, we removed the attP sequence and replaced the single epitope sequences with a 3HA tag in our previous report (36). In the modified vector, we cloned two homologous regions (5’HR and 3’HR) from each SSU gene individually. Details of all cloning steps are available in the Supporting Information.

### 2. Parasite lines, parasite culture, and transfection

D10 wildtype *P. falciparum* was used in this study. Asexual *P. falciparum* parasites were cultured with RPMI-1640 medium supplemented with 0.5% Albumax I (Gibco by ThermoFisher Scientific), sodium bicarbonate (2.1 g/L, Corning by Fisher Scientific), HEPES (15 mM, MilliporeSigma), hypoxanthine (10 mg/L, Fisher Scientific), and gentamycin (50 mg/L, VWR) and maintained as previously described (36). Prior to transfections, template vectors were digested by a specific restriction enzyme overnight to linearize the plasmid (pMG75noP-PfmRPS12-8apt-3HA was digested with *EcoR*V, pMG75noP-PfmRPS17-8apt-3HA and pMG75noP-PfmRPS18-8apt-3HA were done by *Sac*II.) Electroporations were performed in cultures of 5-6% rings added with linearized template plasmids (40 µg per electroporation) together with circular gRNA plasmids (50 µg per gRNA per electroporation). Post electroporation, parasites were cultured with anhydrotetracycline medium at 250 nM (aTc^+^, MilliporeSigma) for 2 days before blasticidin /aTc^+^ medium was used (blasticidin at 2.5 µg/mL, InvivoGen).

### 3. Western blot

Parasitized cultures were treated with saponin (0.05%) in PBS supplemented with a 1x protease inhibitor cocktail (P8215, MilliporeSigma). The parasitized pellet was resuspended in ∼ 10 volumes of 2% SDS / 62 mM Tris-HCl (pH 6.8) and mixed by repeated pipetting. The sample was further mixed on a tube rotator for 4-6 h at room temperature or overnight at 4 °C. The lysate was spun down at top speed for 5 min and the supernatant was used for SDS-PAGE. The blot was blocked and incubated with a mouse monoclonal anti-HA primary antibody (sc-7392, Santa Cruz Biotechnology) at 1:7500 and a HRP conjugated secondary goat anti-mouse antibody (A16078, ThermoFisher Scientific) at 1:10,000. The blot was later probed with a rabbit anti-PfExp2 primary antibody (a gift from Dr. James Burns at Drexel University) at 1:10,000 followed by a HRP conjugated secondary goat anti-rabbit antibody (31460, ThermoFisher Scientific) at 1:20,000. Other steps followed the standard protocol.

### 4. Immunofluorescence assay

Parasite cultures of a 50 µL packed volume were pre-labeled with 60 nM MitoTracker Red CMXRos (Life technologies by ThermoFisher Scientific) for 30 min and washed three times with PBS to remove extra MitoTracker. The cells were fixed with 4% paraformaldehyde/0.0075% glutaraldehyde for 1 h at 37 °C on a rotator. They were washed, permeabilized with 0.5% Triton X-100/PBS for 10 min, treated with 0.1 mg/ml NaBH4 for 5 min, and blocked with 5% BSA overnight. The parasites were incubated with the HA probe (sc-7392, Santa Cruz Biotechnology) at 1:250 for overnight at 4 °C, washed three times with PBS and added with the secondary antibody (Alexa Fluor 488 Conjugated anti-mouse antibody, Life technologies by ThermoFisher Scientific) at 1:250 for overnight at 4 °C. The parasites were washed 3 times with PBS, resuspended in anti-fade buffer (S2828, ThermoFisher Scientific) and visualized under a Leica SP8 confocal microscope. Images were processed using Leica Application Suite X software.

### 5. Immuno-electron microscopy

The D10-PfmRPS18 culture was synchronized with alanine (0.5 M in 10 mM HEPES, pH 7.6) and enriched by a MACS Cell Separation Column (MiltenyiBiotec) at trophozoite stages to reach a high parasitemia (∼ 95%). The enriched parasites in a volume of 25-50 µL were fixed with 1 mL of 2% paraformaldehyde /2.5% glutaraldehyde /100 mM sodium cacodylate at room temperature on a rotator for 1 hr. Samples were washed in sodium cacodylate buffer, embedded in 10% gelatin, and infiltrated overnight with 2.3 M sucrose/20% polyvinyl pyrrolidone in PIPES/MgCl2 at 4 °C. Samples were trimmed, frozen in liquid nitrogen, and sectioned with a Leica Ultracut UCT7 cryo-ultramicrotome (Leica Microsystems Inc.). Ultrathin sections of 50 nm were blocked with 5% FBS (fetal bovine serum)/5% NGS (normal goat serum) for 30 min and subsequently incubated with mouse anti-HA (sc-7392, Santa Cruz Biotechnology) for 1 h at room temperature. Following washes in block buffer, sections were incubated with goat anti-mouse IgG (H+L) 18 nm colloidal gold conjugated secondary antibody (Jackson ImmunoResearch Laboratories) for 1 h. Sections were stained with 0.3% uranyl acetate/2% methyl cellulose and viewed on a JEOL 1200 EX transmission electron microscope (JEOL USA Inc.) equipped with an AMT 8 megapixel digital camera and AMT Image Capture Engine V602 software (Advanced Microscopy Techniques).

All labeling experiments were conducted in parallel with controls omitting the primary antibody. These controls were consistently negative at the concentration of colloidal gold conjugated secondary antibodies used in these studies. Immuno-EM was performed at the Molecular Microbiology Imaging Facility at Washington University in St. Louis, MO.

### 6. Parasites growth curves and knockdown experiment

Parasite cultures were tightly synchronized with several rounds of alanine treatment (0.5 M alanine / 10 mM HEPES, pH 7.6) and trophozoite / schizont stages were isolated using a percoll gradient (89428-524, GE Healthcare Life Sciences). The highly enriched parasites were washed 3 times with regular RPMI (aTc^-^) and inoculated into two new T25 flasks, each containing ∼ 10 µL of parasitized pellet, 1 mL of fresh blood and 10 mL of medium with or without aTc (aTc^+^, aTc^-^). In the next few intraerythrocytic life cycles (IDCs), cultures were split 1:5 every cycle. At each split, 80% of the culture from aTc^+^ and aTc^-^ conditions was used to collect protein samples for Western blot analysis. Thin blood smears were made from both cultures to monitor parasitemia, which was determined from counting 1000 RBCs per smear under a Leica light microscope.

### 7. Mitochondrion preparation (Mito-prep) and *bc*_1_ complex enzymatic activity measurement

For each parasite line (D10-PfmRPS12-3HA, D10-PfmRPS17-3HA or D10-PfmRPS18-3HA), two mito-preps were done to isolate a cellular fraction enriched in parasite mitochondria from parasites grown with or without aTc. With parasites grown in the presence of aTc, each line was tightly synchronized with alanine and expanded to a large volume (8-10 T175 flasks). The saponin treated parasite pellet was disrupted using a nitrogen cavitation at 1000 psi and passed through a MACS CS column (Miltenyi Biotec) to remove the majority of hemozoin. The eluted material from the magnetic column was spun down at 23,000 *g*. This material contains enriched mitochondria or mitochondrial membranes and is suitable for *in vitro bc*_*1*_ enzymatic assay, as shown in our previous studies (36,61). For parasites grown in the absence of aTc, cultures were first grown in aTc^+^ medium to a moderate volume (2-3 T175 flasks) and parasites were isolated by a percoll gradient and inoculated into new T175 flasks supplemented with fresh blood and regular medium (no aTc). The aTc^-^ cultures were maintained for a desired time period and mito-prep was done after removal of aTc for 2 IDCs (D10-PfmRPS17-3HA), 4 IDCs (D10-PfmRPS12-3HA) or 5 IDCs (D10-PfmRPS18-3HA) using the same procedure as described above. For *in vitro bc*_*1*_ enzymatic assay, the cytochrome *c* reductase activity of the *bc*_*1*_ complex was determined using a CLARiTY integrating spectrophotometer (OLIS, Bogart, GA). Briefly, the progress of a 300 µL of *cyt c* reductase reaction, which consisted of mito-prep samples (containing *bc*_*1*_ complex), reduced decylubiquinol (electron donor), oxidized horse heart cytochrome *c* (electron acceptor) and KCN (inhibiting Complex IV activity) in buffer at pH 7.4, was recorded at 550 nm using the spectrophotometer. Other procedures followed our published studies (36,48).

### 8. Drug sensitivity assay (The malaria SYBR Green I-based fluorescence assay, MSF)

Compounds used in this study included DSM-265 (HY-100184, MedCamExpress), atovaquone (A7986, MilliporeSigma), ELQ-300 (synthesized as previously described (24)), proguanil (G7048, MilliporeSigma), artemisinin (a gift from Dr. Jianping Song at Guangzhou University of Chinese Medicine, China), and chloroquine (AC455240250, Fisher Scientific). MSF assays were performed based on published protocols (62,63). Briefly, parasites cultured in aTc^+^ or aTc^-^ medium were exposed to drugs diluted in serial dilutions for 72 h (starting from ring stage parasites). Then the culture was lysed with buffer containing SYBR Green I and examined for relative fluorescence units (RFU) with a Tecan plate reader (Infinite F Plex). Details of setting up the assay and analyzing the data are available in the Supporting Information.

### 9. Metabolic profiling of PfmRPS17 line

The parasite line D10-PfmRPS17-3HA was synchronized and expanded to a moderate volume (2-4 T175 flasks) and late stage parasites (trophozoite/schizont) were isolated using a percoll gradient. The purified parasites were evenly inoculated into two new flasks, added with fresh blood and maintained with or without aTc (time zero). Twenty-four h post time zero, metabolites from aTc^+^ and aTc^-^ cultures were collected every 12 h up to 96 h (in total of 7 time points). The extracted metabolites were analyzed following a published protocol (49). Other details are provided in the Supporting Information.

## Acknowledgements

We are grateful for the active support from members of the Center for Molecular Parasitology in the Department of Microbiology and Immunology at Drexel University College of Medicine. We thank Dr. Josh Beck (Iowa State University, United States), Dr. Jacquin Niles (MIT, United States), Dr. James Burns (Drexel University, United States) and Dr. Jianping Song (Guangzhou University of Chinese Medicine, China) for sharing plasmid vectors and reagents. We thank Dr. Wandy Beatty at the Molecular Microbiology Imaging Facility at Washington University in St. Louis for conducting immune-EM studies.

## Conflict of interest

The authors declare that they have no conflicts of interest with the contents of this article.

## Author contributions

HK and JZ designed the experiments. LL generated transfection plasmids and the parasite lines and performed genotypic and phenotypic analyses with assistance from SW. MM conducted *bc1* assays. JM and ML analyzed metabolite samples by HPLC/mass spectrometry. MWM constructed M-Cas9-yDHOD(-). MKR synthesized ELQ-300 and HDQ. LL and HK wrote the manuscript which was edited by MM, JM, SD, MWM, MKR, ML and JZ.

## FOOTNOTES

* This work was supported by an NIH career transition award (K22, K22AI127702) to Dr. Hangjun Ke and funds from the United States Department of Veterans Affairs, Veterans Health Administration, Office of Research and Development Program Award number i01 BX003312 to M.K.R. M.K.R. is a recipient of a VA Research Career Scientist Award (14S-RCS001). Research reported in this publication was also supported by the US National Institutes of Health under award number AI100569 (M.K.R.).

### The abbreviations used are

PfmRPS12: Plasmodium falciparum mitochondrial ribosomal protein S12
PfmRPS17: Plasmodium falciparum mitochondrial ribosomal protein S17
PfmRPS18: Plasmodium falciparum mitochondrial ribosomal protein S18
mitoribosomes: mitochondrial ribosome
mtETC: mitochondrial electron transport chain
mtDNA: mitochondrial DNA
cyt: cytochrome
cox1: cytochrome oxidase subunit I
cox3: cytochrome oxidase subunit III
TetR: tetracycline repressor
DOZI: development of zygote inhibited
cryo-EM: cryoelectron microscopy
gRNA: guider RNA
aTc: anhydrotetracycline
DHOD: dihydroorotate dehydrogenase
Qd: decylubiquinone
SSU: small subunit
LSU: large subunit
ELQ-300: endochin like quinolone-300
HDQ: hydroxy-2-dodecyl-4-(1H)quinolone

## REFERENCES

1. WHO. (2016) World Malaria Report.

2. WHO. (2017) World Malaria Report.

3. WHO. (2018) World Malaria Report.

4. Gardner, M. J., Hall, N., Fung, E., White, O., Berriman, M., Hyman, R. W., Carlton, J. M., Pain, A., Nelson, K. E., Bowman, S., Paulsen, I. T., James, K., Eisen, J. A., Rutherford, K., Salzberg, S. L., Craig, A., Kyes, S., Chan, M. S., Nene, V., Shallom, S. J., Suh, B., Peterson, J., Angiuoli, S., Pertea, M., Allen, J., Selengut, J., Haft, D., Mather, M. W., Vaidya, A. B., Martin, D. M., Fairlamb, A. H., Fraunholz, M. J., Roos, D. S., Ralph, S. A., McFadden, G. I., Cummings, L. M., Subramanian, G. M., Mungall, C., Venter, J. C., Carucci, D. J., Hoffman, S. L., Newbold, C., Davis, R. W., Fraser, C. M., and Barrell, B. (2002) Genome sequence of the human malaria parasite Plasmodium falciparum. Nature 419, 498–511

5. Wong, W., Bai, X. C., Brown, A., Fernandez, I. S., Hanssen, E., Condron, M., Tan, Y. H., Baum, J., and Scheres, S. H. (2014) Cryo-EM structure of the Plasmodium falciparum 80S ribosome bound to the anti-protozoan drug emetine. Elife 3

6. Sun, M., Li, W., Blomqvist, K., Das, S., Hashem, Y., Dvorin, J. D., and Frank, J. (2015) Dynamical features of the Plasmodium falciparum ribosome during translation. Nucleic Acids Res 43, 10515–10524

7. Habib, S., Vaishya, S., and Gupta, K. (2016) Translation in Organelles of Apicomplexan Parasites. Trends Parasitol 32, 939–952

8. Gupta, A., Shah, P., Haider, A., Gupta, K., Siddiqi, M. I., Ralph, S. A., and Habib, S. (2014) Reduced ribosomes of the apicoplast and mitochondrion of Plasmodium spp. and predicted interactions with antibiotics. Open Biol 4, 140045

9. Dahl, E. L., and Rosenthal, P. J. (2007) Multiple antibiotics exert delayed effects against the Plasmodium falciparum apicoplast. Antimicrob Agents Chemother 51, 3485–3490

10. Goodman, C. D., Su, V., and McFadden, G. I. (2007) The effects of anti-bacterials on the malaria parasite Plasmodium falciparum. Mol Biochem Parasitol 152, 181–191

11. Yeh, E., and DeRisi, J. L. (2011) Chemical rescue of malaria parasites lacking an apicoplast defines organelle function in blood-stage Plasmodium falciparum. PLoS Biol 9, e1001138

12. Schon, E. A., DiMauro, S., and Hirano, M. (2012) Human mitochondrial DNA: roles of inherited and somatic mutations. Nat Rev Genet 13, 878–890

13. Hikosaka, K., Kita, K., and Tanabe, K. (2013) Diversity of mitochondrial genome structure in the phylum Apicomplexa. Mol Biochem Parasitol 188, 26–33

14. Vaidya, A. B., Akella, R., and Suplick, K. (1989) Sequences similar to genes for two mitochondrial proteins and portions of ribosomal RNA in tandemly arrayed 6-kilobase-pair DNA of a malarial parasite. Mol Biochem Parasitol 35, 97–107

15. Suplick, K., Akella, R., Saul, A., and Vaidya, A. B. (1988) Molecular cloning and partial sequence of a 5.8 kilobase pair repetitive DNA from Plasmodium falciparum. Mol Biochem Parasitol 30, 289–290

16. Gray, M. W., Lang, B. F., and Burger, G. (2004) Mitochondria of protists. Annu Rev Genet 38, 477–524

17. Feagin, J. E., Harrell, M. I., Lee, J. C., Coe, K. J., Sands, B. H., Cannone, J. J., Tami, G., Schnare, M. N., and Gutell, R. R. (2012) The fragmented mitochondrial ribosomal RNAs of Plasmodium falciparum. PLoS One 7, e38320

18. Srivastava, I. K., Rottenberg, H., and Vaidya, A. B. (1997) Atovaquone, a broad spectrum antiparasitic drug, collapses mitochondrial membrane potential in a malarial parasite. J Biol Chem 272, 3961–3966

19. Vaidya, A. B., and Mather, M. W. (2000) Atovaquone resistance in malaria parasites. Drug Resist Updat 3, 283–287

20. Fry, M., and Pudney, M. (1992) Site of action of the antimalarial hydroxynaphthoquinone, 2- [trans-4-(4’-chlorophenyl) cyclohexyl]-3-hydroxy-1,4-naphthoquinone (566C80). Biochem Pharmacol 43, 1545–1553

21. Ke, H., & Mather, M. W. (2017) Targeting mitochondrial functions as antimalarial regime, what is next?. Current Clinical Microbiology 4, 175–191

22. Frueh, L., Li, Y., Mather, M. W., Li, Q., Pou, S., Nilsen, A., Winter, R. W., Forquer, I. P., Pershing, A. M., Xie, L. H., Smilkstein, M. J., Caridha, D., Koop, D. R., Campbell, R. F., Sciotti, R. J., Kreishman-Deitrick, M., Kelly, J. X., Vesely, B., Vaidya, A. B., and Riscoe, M. K. (2017) Alkoxycarbonate Ester Prodrugs of Preclinical Drug Candidate ELQ-300 for Prophylaxis and Treatment of Malaria. ACS Infect Dis 3, 728–735

23. Stickles, A. M., Smilkstein, M. J., Morrisey, J. M., Li, Y., Forquer, I. P., Kelly, J. X., Pou, S., Winter, R. W., Nilsen, A., Vaidya, A. B., and Riscoe, M. K. (2016) Atovaquone and ELQ-300 Combination Therapy as a Novel Dual-Site Cytochrome bc1 Inhibition Strategy for Malaria. Antimicrob Agents Chemother 60, 4853–4859

24. Miley, G. P., Pou, S., Winter, R., Nilsen, A., Li, Y., Kelly, J. X., Stickles, A. M., Mather, M. W., Forquer, I. P., Pershing, A. M., White, K., Shackleford, D., Saunders, J., Chen, G., Ting, L. M., Kim, K., Zakharov, L. N., Donini, C., Burrows, J. N., Vaidya, A. B., Charman, S. A., and Riscoe, M. K. (2015) ELQ-300 prodrugs for enhanced delivery and single-dose cure of malaria. Antimicrob Agents Chemother 59, 5555–5560

25. Brunk, C. F., Lee, L. C., Tran, A. B., and Li, J. (2003) Complete sequence of the mitochondrial genome of Tetrahymena thermophila and comparative methods for identifying highly divergent genes. Nucleic Acids Res 31, 1673–1682

26. Boer, P. H., and Gray, M. W. (1988) Transfer RNA genes and the genetic code in Chlamydomonas reinhardtii mitochondria. Curr Genet 14, 583–590

27. Amunts, A., Brown, A., Bai, X. C., Llacer, J. L., Hussain, T., Emsley, P., Long, F., Murshudov, G., Scheres, S. H. W., and Ramakrishnan, V. (2014) Structure of the yeast mitochondrial large ribosomal subunit. Science 343, 1485–1489

28. Desai, N., Brown, A., Amunts, A., and Ramakrishnan, V. (2017) The structure of the yeast mitochondrial ribosome. Science 355, 528–531

29. Brown, A., Amunts, A., Bai, X. C., Sugimoto, Y., Edwards, P. C., Murshudov, G., Scheres, S. H. W., and Ramakrishnan, V. (2014) Structure of the large ribosomal subunit from human mitochondria. Science 346, 718–722

30. Amunts, A., Brown, A., Toots, J., Scheres, S. H. W., and Ramakrishnan, V. (2015) Ribosome. The structure of the human mitochondrial ribosome. Science 348, 95–98

31. Brown, A., Rathore, S., Kimanius, D., Aibara, S., Bai, X. C., Rorbach, J., Amunts, A., and Ramakrishnan, V. (2017) Structures of the human mitochondrial ribosome in native states of assembly. Nat Struct Mol Biol 24, 866–869

32. Greber, B. J., Boehringer, D., Leitner, A., Bieri, P., Voigts-Hoffmann, F., Erzberger, J. P., Leibundgut, M., Aebersold, R., and Ban, N. (2014) Architecture of the large subunit of the mammalian mitochondrial ribosome. Nature 505, 515–519

33. Greber, B. J., Boehringer, D., Leibundgut, M., Bieri, P., Leitner, A., Schmitz, N., Aebersold, R., and Ban, N. (2014) The complete structure of the large subunit of the mammalian mitochondrial ribosome. Nature 515, 283–286

34. Greber, B. J., Bieri, P., Leibundgut, M., Leitner, A., Aebersold, R., Boehringer, D., and Ban, N. (2015) Ribosome. The complete structure of the 55S mammalian mitochondrial ribosome. Science 348, 303–308

35. Ramrath, D. J. F., Niemann, M., Leibundgut, M., Bieri, P., Prange, C., Horn, E. K., Leitner, A., Boehringer, D., Schneider, A., and Ban, N. (2018) Evolutionary shift toward protein-based architecture in trypanosomal mitochondrial ribosomes. Science 362

36. Ke, H., Dass, S., Morrisey, J. M., Mather, M. W., and Vaidya, A. B. (2018) The mitochondrial ribosomal protein L13 is critical for the structural and functional integrity of the mitochondrion in Plasmodium falciparum. J Biol Chem 293, 8128–8137

37. Ganesan, S. M., Falla, A., Goldfless, S. J., Nasamu, A. S., and Niles, J. C. (2016) Synthetic RNA-protein modules integrated with native translation mechanisms to control gene expression in malaria parasites. Nat Commun 7, 10727

38. Lacombe, A., Maclean, A. E., Ovciarikova, J., Tottey, J., Muhleip, A., Fernandes, P., and Sheiner, L. (2019) Identification of the Toxoplasma gondii mitochondrial ribosome, and characterisation of a protein essential for mitochondrial translation. Mol Microbiol 112, 1235–1252

39. Shajani, Z., Sykes, M. T., and Williamson, J. R. (2011) Assembly of bacterial ribosomes. Annu Rev Biochem 80, 501–526

40. Englmeier, R., Pfeffer, S., and Forster, F. (2017) Structure of the Human Mitochondrial Ribosome Studied In Situ by Cryoelectron Tomography. Structure 25, 1574–1581 e1572

41. Pease, B. N., Huttlin, E. L., Jedrychowski, M. P., Talevich, E., Harmon, J., Dillman, T., Kannan, N., Doerig, C., Chakrabarti, R., Gygi, S. P., and Chakrabarti, D. (2013) Global analysis of protein expression and phosphorylation of three stages of Plasmodium falciparum intraerythrocytic development. J Proteome Res 12, 4028–4045

42. Ramaswamy, P., and Woodson, S. A. (2009) Global stabilization of rRNA structure by ribosomal proteins S4, S17, and S20. J Mol Biol 392, 666–677

43. Vila-Sanjurjo, A., Lu, Y., Aragonez, J. L., Starkweather, R. E., Sasikumar, M., and O’Connor, M. (2007) Modulation of 16S rRNA function by ribosomal protein S12. Biochim Biophys Acta 1769, 462–471

44. Zhang, M., Wang, C., Otto, T. D., Oberstaller, J., Liao, X., Adapa, S. R., Udenze, K., Bronner, I. F., Casandra, D., Mayho, M., Brown, J., Li, S., Swanson, J., Rayner, J. C., Jiang, R. H. Y., and Adams, J. H. (2018) Uncovering the essential genes of the human malaria parasite Plasmodium falciparum by saturation mutagenesis. Science 360

45. Murphy, S. C., Duke, E. R., Shipman, K. J., Jensen, R. L., Fong, Y., Ferguson, S., Janes, H. E., Gillespie, K., Seilie, A. M., Hanron, A. E., Rinn, L., Fishbaugher, M., VonGoedert, T., Fritzen, E., Kappe, S. H., Chang, M., Sousa, J. C., Marcsisin, S. R., Chalon, S., Duparc, S., Kerr, N., Mohrle, J. J., Andenmatten, N., Rueckle, T., and Kublin, J. G. (2018) A Randomized Trial Evaluating the Prophylactic Activity of DSM265 Against Preerythrocytic Plasmodium falciparum Infection During Controlled Human Malarial Infection by Mosquito Bites and Direct Venous Inoculation. J Infect Dis 217, 693–702

46. Painter, H. J., Morrisey, J. M., Mather, M. W., and Vaidya, A. B. (2007) Specific role of mitochondrial electron transport in blood-stage Plasmodium falciparum. Nature 446, 88–91

47. Lane, K. D., Mu, J., Lu, J., Windle, S. T., Liu, A., Sun, P. D., and Wellems, T. E. (2018) Selection of Plasmodium falciparum cytochrome B mutants by putative PfNDH2 inhibitors. Proc Natl Acad Sci U S A 115, 6285–6290

48. Ke, H., Ganesan, S. M., Dass, S., Morrisey, J. M., Pou, S., Nilsen, A., Riscoe, M. K., Mather, M. W., and Vaidya, A. B. (2019) Mitochondrial type II NADH dehydrogenase of Plasmodium falciparum (PfNDH2) is dispensable in the asexual blood stages. PLoS One 14, e0214023

49. Allman, E. L., Painter, H. J., Samra, J., Carrasquilla, M., and Llinas, M. (2016) Metabolomic Profiling of the Malaria Box Reveals Antimalarial Target Pathways. Antimicrob Agents Chemother 60, 6635–6649

50. Karnkowska, A., Vacek, V., Zubacova, Z., Treitli, S. C., Petrzelkova, R., Eme, L., Novak, L., Zarsky, V., Barlow, L. D., Herman, E. K., Soukal, P., Hroudova, M., Dolezal, P., Stairs, C. W., Roger, A. J., Elias, M., Dacks, J. B., Vlcek, C., and Hampl, V. (2016) A Eukaryote without a Mitochondrial Organelle. Curr Biol 26, 1274–1284

51. Sturm, A., Mollard, V., Cozijnsen, A., Goodman, C. D., and McFadden, G. I. (2015) Mitochondrial ATP synthase is dispensable in blood-stage Plasmodium berghei rodent malaria but essential in the mosquito phase. Proc Natl Acad Sci U S A 112, 10216–10223

52. Bushell, E., Gomes, A. R., Sanderson, T., Anar, B., Girling, G., Herd, C., Metcalf, T., Modrzynska, K., Schwach, F., Martin, R. E., Mather, M. W., McFadden, G. I., Parts, L., Rutledge, G. G., Vaidya, A. B., Wengelnik, K., Rayner, J. C., and Billker, O. (2017) Functional Profiling of a Plasmodium Genome Reveals an Abundance of Essential Genes. Cell 170, 260–272 e268

53. Zheng, W., Zhang, C., Bell, E. W., and Zhang, Y. (2019) I-TASSER gateway: A protein structure and function prediction server powered by XSEDE. Future Gener Comput Syst 99, 73–85

54. Moller-Hergt, B. V., Carlstrom, A., Stephan, K., Imhof, A., and Ott, M. (2018) The ribosome receptors Mrx15 and Mba1 jointly organize cotranslational insertion and protein biogenesis in mitochondria. Mol Biol Cell 29, 2386–2396

55. Rouault, T. A. (2012) Biogenesis of iron-sulfur clusters in mammalian cells: new insights and relevance to human disease. Dis Model Mech 5, 155–164

56. Rouault, T. A., and Tong, W. H. (2005) Iron-sulphur cluster biogenesis and mitochondrial iron homeostasis. Nat Rev Mol Cell Biol 6, 345–351

57. Xu, P., Widmer, G., Wang, Y., Ozaki, L. S., Alves, J. M., Serrano, M. G., Puiu, D., Manque, P., Akiyoshi, D., Mackey, A. J., Pearson, W. R., Dear, P. H., Bankier, A. T., Peterson, D. L., Abrahamsen, M. S., Kapur, V., Tzipori, S., and Buck, G. A. (2004) The genome of Cryptosporidium hominis. Nature 431, 1107–1112

58. Spillman, N. J., Beck, J. R., Ganesan, S. M., Niles, J. C., and Goldberg, D. E. (2017) The chaperonin TRiC forms an oligomeric complex in the malaria parasite cytosol. Cell Microbiol 19

59. Chen, B., Gilbert, L. A., Cimini, B. A., Schnitzbauer, J., Zhang, W., Li, G. W., Park, J., Blackburn, E. H., Weissman, J. S., Qi, L. S., and Huang, B. (2013) Dynamic imaging of genomic loci in living human cells by an optimized CRISPR/Cas system. Cell 155, 1479–1491

60. Dang, Y., Jia, G., Choi, J., Ma, H., Anaya, E., Ye, C., Shankar, P., and Wu, H. (2015) Optimizing sgRNA structure to improve CRISPR-Cas9 knockout efficiency. Genome Biol 16, 280

61. Mather, M. W., Morrisey, J. M., and Vaidya, A. B. (2010) Hemozoin-free Plasmodium falciparum mitochondria for physiological and drug susceptibility studies. Mol Biochem Parasitol 174, 150–153

62. Smilkstein, M., Sriwilaijaroen, N., Kelly, J. X., Wilairat, P., and Riscoe, M. (2004) Simple and inexpensive fluorescence-based technique for high-throughput antimalarial drug screening. Antimicrob Agents Chemother 48, 1803–1806

63. Johnson, J. D., Dennull, R. A., Gerena, L., Lopez-Sanchez, M., Roncal, N. E., and Waters, N. C. (2007) Assessment and continued validation of the malaria SYBR green I-based fluorescence assay for use in malaria drug screening. Antimicrob Agents Chemother 51, 1926–1933

